# SPONGE: Simple Prior Omics Network GEnerator

**DOI:** 10.1101/2024.10.24.619989

**Authors:** Ladislav Hovan, Marieke L. Kuijjer

## Abstract

Gene regulatory networks modelled from experimental data can be improved through the use of prior biological knowledge on e.g. transcription factor binding. There are several tools that utilise this information. However, the prior networks used with them are often not updated and may fail to reflect the most up-to-date information. Here we present SPONGE, a Python module designed to access information across biological databases, chiefly JASPAR and STRING, to model two types of networks—a prior gene regulatory network mapping transcription factors to genes based on their predicted binding sites, and a prior protein-protein interaction network mapping potential interactions between transcription factors. SPONGE is mainly designed to work with the PANDA algorithm and the corresponding NetZoo family of tools. However, the networks are provided in an easily adaptable format for other tools. SPONGE was designed with ease of use in mind, and it provides sensible default values for all of its parameters while giving the users freedom to fine-tune them. The code for the Python module and the documentation can be found in [our GitHub repository](https://github.com/ladislav-hovan/sponge).

## 1. Introduction

The process of transcriptional regulation in humans is complex and still a subject of intensive research Kim and Wysocka (2023). It is, however, already known that this process is crucial to our understanding of many diseases, including cancer Ankill et al. (2022). One way to model this process is through gene regulatory network (GRN) inference Zitnik et al. (2024). GRNs are networks that describe the regulation of gene expression with edges, representing regulatory interactions, drawn between regulators, such as transcription factors (TFs) and their target genes.

Together with our collaborators, we have created a large repository of tools for creating and analysing GRNs called NetZoo Guebila et al. (2023). One of the central tools for GRN reconstruction in NetZoo is called PANDA (Passing Attributes between Networks for Data Assimilation) Glass et al. (2013); van IJzendoorn et al. (2016). This approach utilises three main sources of information: a prior network of potential gene regulation based on binding sites of TFs in gene promoters, a prior network of potential protein-protein interaction (PPI) between TFs, and gene expression profiles measured for multiple samples, which is used to calculate a gene co-expression network. The main idea behind PANDA is that utilising our prior knowledge of gene regulation through the two prior networks results in a more robust and reliable GRN than relying entirely on the experimental data, which may fail to capture gene regulation directly Chen et al. (2014); McCalla et al. (2023). Therefore, the quality of these prior networks is as important as that of the experiments.

In order to facilitate future creation of prior GRNs a PPI networks, we developed a prior network generator, which we call SPONGE (Simple Prior Omics Network GEnerator), as a dedicated contribution to the NetZoo. The main motivation for creating SPONGE is twofold: ease of update and reproducibility. The databases that provide data which are used to generate prior networks are updated relatively frequently. However, in our experience, once a prior network has been provided, it tends to circulate between users and research groups for a long time without updates. We believe this to be a result of the perceived difficulty in creating prior networks, and we hope that with its user-friendliness, SPONGE will allow people to keep their prior networks as up-to-date as possible.

The second reason is facilitating the reproducibility of prior network creation. Many different ways of constructing prior GRNs exist: various tools to perform motif scans (FIMO Grant et al. (2011), MotifScan Sun et al. (2018), etc.), as well as several hyperparameters to decide on when constructing the prior network. We aim to eliminate some of the possible sources of variation by using the reliable, curated JASPAR Rauluseviciute et al. (2024) database as the source of motifs. The authors of the database also provide the results of motif scans for the entire genome, which allows us to eliminate this time-consuming step. The values of hyperparameters, such as score thresholds and promoter boundaries, are easy to adjust, and the default values (see below) are chosen in a way that is consistent with current practice. Thus, we believe that this tool will make creation of prior networks tailored to one’s preferences easy for everyone. We provide the networks generated with the default settings in September 2024 for anyone to use. They can be found on Zenodo.

## 2. Tool description

SPONGE is a Python module that greatly simplifies the generation of prior GRNs and PPI networks between the TFs present. The data is mainly retrieved from the JASPAR Rauluseviciute et al. (2024) database of TF binding motifs and the STRING Szklarczyk et al. (2023) database of PPIs. Information about human homologs (see below) of vertebrate transcription factors is retrieved from NCBI Datasets O’Leary et al. (2024). Mappings between gene identifiers from UniProt Bateman et al. (2023) are queried at several points. Ensembl Harrison et al. (2024) is also utilised by default in order to get a list of all human transcripts and their respective genes, but its output can be fully replaced by a user-provided file. Currently, SPONGE is designed to work with the human genome only, as to our knowledge most of the PANDA user base works with human networks. However, in the future we plan to expand this to other organisms represented in JASPAR. The schematic of the workflow is described in Fig. 1A.

**Fig 1:**
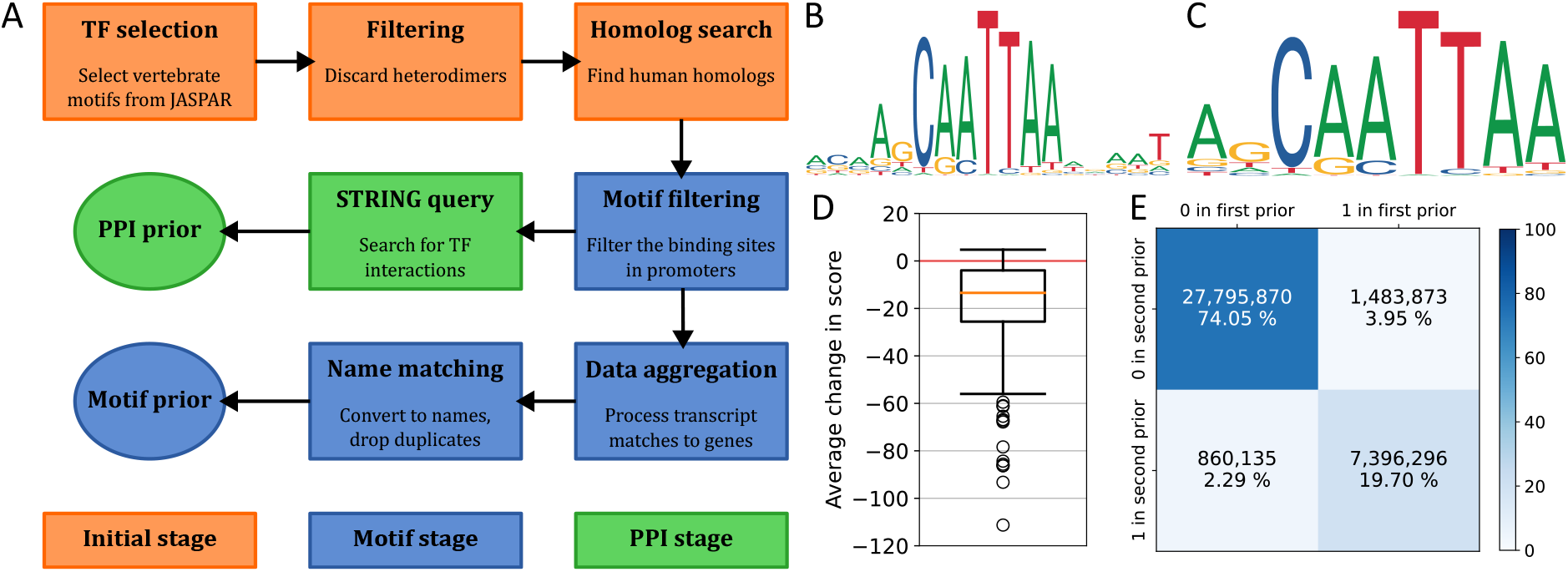
A. Overview of the SPONGE pipeline, divided into three main parts. B. Motif logo for the TF Hmx3 in JASPAR2022 (Motif ID MA0898.1). C. Motif logo for the TF Hmx3 in JASPAR2024 (Motif ID MA0898.2). D. The average change in score for newest versions of TF motifs going from JASPAR2022 to JASPAR2024. Due to the very large number of predicted TF binding sites in the human genome, the analysis visualized here is restricted to the binding sites on chromosome 19. E. The confusion matrix for edges in the prior GRNs generated from JASPAR2022 and JASPAR2024. It is restricted to TFs present in both networks.

As the first thing after initialisation, SPONGE makes sure that it has all the files required for the successful execution of the pipeline. This does not mean that these files have to be provided by the user—SPONGE will attempt to download all the files that are required but not provided. In the default settings, these would be the promoter file listing the genome regions of interest, the Ensembl file mapping transcripts to genes (by default generated at the same time as the promoter file), and finally the bigbed file from JASPAR with all the detected TF binding sites in the genome. The general purpose of these files is to filter out the TF binding sites that occur in promoters, and then link these promoters to their genes. The bigbed file is very large (more than 160 GB in JASPAR2024 for hg38), therefore as an alternative, it is possible to download the files for individual TFs instead as they are being processed. The largest one of these is just under 2 GB in size. However, processing the data this way takes more time (see below). By default, SPONGE keeps all the downloaded files in a temporary directory and will not delete them unless instructed to. In future runs, it will use these files again if not instructed otherwise. Users are free to substitute these files with their own, as long as the required fields are present. The specifications for the files are described in the GitHub repository.

The next step in SPONGE is the initialisation phase (Fig. 1A, highlighted in orange). The newest versions of all the vertebrate motifs are retrieved from JASPAR, followed by optional filtering of heterodimers. Finally, SPONGE uses the data from NCBI Datasets to try to find human homologs of vertebrate TFs. We include this step to increase the number of TFs in the priors, with the justification that the TF motifs are often very highly conserved Nitta et al. (2015). Where these correspond to ones already present in JASPAR in their human version, the human version is preferred. If no homolog is found, the vertebrate motif is discarded.

The next step is the motif stage (Fig. 1A, highlighted in blue). First, SPONGE finds the overlap between the defined regions of interest and the TF binding sites as retrieved from JASPAR. The default definition of a region of interest is from 750 bases upstream to 250 downstream of the transcription start site for all the genes. Motif filtering is the most important part of the entire SPONGE pipeline and it is designed in a way that allows for parallel processing. The actual implementation differs slightly based on whether the JASPAR bigbed file is present. If it is, the processing of each individual chromosome is parallelised and all the TFs are dealt with at the same time. This means that every region of interest only has to be processed once. In case of the on-the-fly download of individual TF tracks, every region of interest is processed once per TF, which inevitably leads to slowdown. However, the processing of individual chromosomes is parallelised. The actual slowdown depends on whether download times are included. If they are, the runtime for bibged processing in our case was around one hour, compared to six hours for on the fly processing. This should only be taken as an illustration, as the actual comparison will depend greatly on the internet connection speed and the number of cores assigned to the algorithm. Only the match with the best score is retained for every combination of region of interest and TF. Also, a minimum score threshold is employed (400 by default).

The result of the motif filtering is the mapping of TFs to transcripts. This mapping is then postprocessed first by aggregating transcript matches into genes, and then optionally converting the gene IDs into names. In case of multiple gene IDs with the same name, only the one with the highest number of TF binding sites is retained. While this may potentially cause problems, the number of genes affected by this is negligible, and it can be avoided entirely by using the gene IDs throughout. Finally, the motif prior is saved as a file in PANDA-compatible format.

At last, the PPI stage (Fig. 1A, highlighted in green) is executed. The TFs which have been found to have binding sites in promoters are queried in the STRING database for their interactions, and these are then saved into a file.

In order to ensure reproducibility, SPONGE keeps track of the versions and retrieval times of all the files it downloads. These fingerprints can be shown, and will be kept track of even after the downloaded files are reused in future runs. This is done through a hidden file in the temporary directory that SPONGE uses. However, SPONGE cannot maintain fingerprints for user-provided files.

As an example of why one may want to update their prior networks as databases are updated, we have run SPONGE for both the newest version of the JASPAR database (JASPAR2024 Rauluseviciute et al. (2024)) and the previous one (JASPAR2022 Castro-Mondragon et al. (2022)). Apart from the usual addition of new TF motifs into the database, one change that has been made was the trimming of TF motifs. Some of the motifs which had bases with little information content near the edges had these less-informative bases removed in the database update. An example of this process for the TF HMX3 (Hmx3 from Mus Musculus, JASPAR motif ID MA0898) can be seen in Fig. 1B and C. As a consequence, the scores calculated are also affected, and in general they are lower in the newer version of JASPAR, as shown in Fig. 1D. This has resulted in an unexpected situation where even though the overall number of TFs has increased in the newer version of JASPAR, the number of TFs in the resulting motif prior was lower (625 vs 672), because some have fallen below the default score threshold. However, for those TFs present in both versions of the motif prior, the results are, not unexpectedly, very similar, with only about 6% of edges being different (Fig. 1E).

## 3. Availability and implementation

SPONGE can be found on GitHub. The repository also contains links to documentation and installation instructions. The most recent release can also be installed via pip and a Docker container is also available. Finally, the repository has been archived on Zenodo to ensure future availability. The prior networks generated with the latest settings can also be found there.

## 4. Conclusion

To conclude, SPONGE provides an easy way to generate prior GRNs and PPI networks and keep up to date with ever improving JASPAR and STRING databases. Although it has mainly been designed with PANDA in mind, these networks can be useful for other applications too. SPONGE can also be utilised differently, for example by replacing the promoter regions with other regions of interest such as enhancers.

In the future, we would like to expand SPONGE with an ability to generate GRNs for organisms other than humans. Although we believe it to be reasonably efficient, performance improvements will also be considered.

## Acknowledgments

The authors would like to acknowledge Katalin Ferenc and Vipin Kumar for their suggestions to make the user interface more usable and intuitive, and Ieva Rauluseviciute for her insights into the JASPAR database. We would also like to acknowledge the contributions from all the other members of the Kuijjer and Mathelier groups at NCMM, especially those who have provided their feedback during our code review meetings.

## Author contributions

L.H. designed, programmed and tested SPONGE. M.L.K. provided supervision. L.H and M.L.K. wrote and edited the manuscript.

## Conflict of interest

None declared.

## Funding

The authors were supported by funding from the Research Council of Norway [187615], Helse Sør-Øst, and the University of Oslo through the Centre for Molecular Medicine Norway (NCMM); the Norwegian Cancer Society [214871, 273592], the Research Council of Norway [313932], and the European Union’s Horizon 2020 research and innovation program under the Marie Sklodowska-Curie grant agreement [801133] (to L.H.).

## Data availability

By default, SPONGE uses exclusively publicly available data from the JASPAR Rauluseviciute et al. (2024), STRING Szklarczyk et al. (2023), NCBI Datasets O’Leary et al. (2024), UniProt Bateman et al. (2023), and Ensembl Harrison et al. (2024) databases. All the code for SPONGE and the code required to reproduce the figures in this manuscript can be found in the GitHub repository.

## Notes

https://github.com/ladislav-hovan/sponge

https://zenodo.org/records/13628784

### Competing Interest Statement

The authors have declared no competing interest.

